# Unveiling the Hidden: Dissecting Liraglutide Oligomerization Dual Pathways via Direct Mass Technology, Electron-Capture Dissociation, and Molecular Dynamics

**DOI:** 10.1101/2025.02.27.640645

**Authors:** Syuan-Ting Kuo, Zhenyu Xi, Xiao Cong, Xin Yan, David H. Russell

## Abstract

Peptide therapeutics have revolutionized drug design strategies, yet the inherent structural flexibility and conjugated moieties of drug molecules present challenges in discovery, rational design, and manufacturing. Liraglutide, a GLP-1 receptor agonist conjugated with palmitic acid at its lysine residue, exemplifies these challenges by forming oligomers, which may compromise efficacy through progressive formation of aggregates. Here, we incorporate native mass spectrometry platforms including electron-capture dissociation (ECD), direct mass technology (DMT), and molecular dynamics (MD) to capture the early oligomerization process of liraglutide. Our findings reveal a restricted C-terminal region upon oligomer formation, as indicated by the reduced release of *z*-ions in ECD analysis. Additionally, we identified the formation of higher-order oligomers (*n*=25–62) by DMT, primarily stabilized by hydrophilic interactions involving preformed stable oligomers (*n*=14). Together, these integrative mass spectrometry results delineate a dual-pathway oligomerization process for liraglutide, demonstrating the power of mass spectrometry in uncovering hidden pathways of self-association. This approach underscores the potential of mass spectrometry as a key tool in the rational design and optimization of peptide-based therapeutics.

## INTRODUCTION

Liraglutide, a glucagon-like peptide-1 receptor agonist (GLP-1 RA) modified through the conjugation of a C16 fatty acid to Lys26, exhibits high efficacy in weight control and the treatment of type II diabetes.^1-4^ This lipidation promotes self-association, which prolongs the half-life of liraglutide in blood circulation by allowing its slow release from oligomers and prevention of renal clearance.^5-7^ To ensure its reliability, it is pivotal to understand how its oligomerization is modulated by physiological conditions such as temperature, ionic strength, and acidity of the solution. Traditional solution-based methods such as ultracentrifugation (AUC), small-angle X-ray scattering (SAXS), and size exclusion chromatography coupled with multi-angle static light scattering (SEC-MALS) have been widely adopted to determine its oligomerization states.^6, 8-11^ These studies consistently show that liraglutide forms oligomers *n*=12-14 in mildly acid conditions (pH<6.8) and oligomers *n*=6-8 in neutral or slightly basic conditions (pH>7.0), with the oligomeric transition being reversible and sensitive to pH alternations.^6^

While conventional methods are robust and widely adopted, they primarily provide ensemble-averaged measurements of oligomers in solution.^12^ This averaging can oversimplify analyte properties by emphasizing dominant oligomeric species while overlooking low-abundance intermediates, which may trigger unexpected cascading on- and off-target aggregation. For example, in neurodegenerative diseases, low-abundance pre-fibrillar oligomers are increasingly recognized as critical predictors of disease progression, with their modulation influencing the final fibrillation morphology.^13^ Similarly, studies on liraglutide have explored the relationship between soluble oligomers and final aggregate morphology.^6^

However, the presence of soluble oligomers failed to predict the kinetics and morphology of final insoluble products in current studies. This may indicate the existence of unresolved intermediate or pre-fibrillar oligomers that remain undefined by conventional techniques. An advanced analytical platform capable of resolving heterogeneous and low-abundance oligomers is therefore necessary to fully elucidate oligomerization of liraglutide.

Among current analytical platforms, native mass spectrometry (nMS) has emerged as a powerful tool for resolving highly heterogeneous systems.^14-16^ Pioneering high resolution native MS platforms have resolved minute differences in masses of small molecules (i.e. lipid and metal ions) bound to membrane proteins.^17-19^ Combined with a variable temperature electrospray ionization (vT-ESI) device, high mass resolution enables the simultaneous determination of thermodynamic signatures (ΔG, ΔH, and TΔS) of up to 14 ATP binding events to the 801 kDa GroEL to study the effects of solution composition (i.e. temperature, buffer, and cofactors) in a highly heterogeneous system.^20-22^ Beyond stoichiometric information, higher-order structural information can be extracted through advanced dissociation techniques. For example, comparing c- and z-ions released from electron-capture dissociation (ECD) allows the identification of metal-binding sites, weakly bound ligands, structural modulations of proteins, and multimerization interfaces of protein complexes. ^23-26^ These capabilities make nMS a unique method for studying complex biological assemblies that are difficult to resolve using solution-based techniques.

Recent advance in charge detection mass spectrometry (CDMS) and Direct Mass Technology (DMT) have further transformed mass analysis by enabling charge assignment for individual ions without relying on isotope patterns or multiple charge states.^27-29^ This advancement improves the detection of low-abundance and heterogeneous species with overlapping *m/z* signals, making it particularly useful for analyzing large biomolecules such as antibodies,^30^ spike proteins,^31^ and adeno-associated viruses (AAVs).^32-34^ These methods have been further investigated to resolve oligomer identities and structures to understand oligomerization mechanisms. In the analysis of monoclonal antibody (mAb), both compact and elongated oligomers with identical masses have been differentiated by two distinct charge distributions. A structural rearrangement was rationalized in pentameric and decameric mAbs, evidenced by their high abundance of compact conformers and necessitated the need for resolving conformational heterogeneity by charges.^35^ This charge-conformer relationship has been further developed as a high-throughput platform to characterize oligomers under stress conditions, resolving unusual bovine serum albumin (BSA) oligomers consisting of up to 225 monomers.^36^

Here, we present integrative mass spectrometry techniques to characterize liraglutide oligomers. ECD differentiated a restricted dynamic upon liraglutide oligomerization, which is necessary for growth to higher molecular weight (HMW) oligomers. Previously undetected oligomers of liraglutide were resolved by direct mass technology (DMT). These experimental findings were corroborated by molecular dynamics simulations, providing explanatory mechanisms of the unusual discontinuity of oligomeric states observed in the mass domain. We established a framework of using mass spectrometry to study multiple levels in the stage of oligomerization and dissected the oligomerization pathways utilizing both hydrophobic and hydrophilic residues in the GLP-1 RA.

## METHODS

## MATERIALS

Liraglutide was purchased from MedChemExpress LLC (Monmouth Junction, NJ) with 98.91% purity. Liraglutide was prepared and analyzed by reconstituting the powder in 20 mM ammonium acetate to reach the concentration of 1 mg/mL without any pretreatment. The pH was adjusted by either ammonium hydroxide or acetic acid to desired value after dissolution. The incubation temperature was 37°C unless otherwise specified.

### CHARACTERIZATION OF LIRAGLUTIDE OLIGOMERS BY NATIVE MASS SPECTROMETRY

Characterization of liraglutide oligomers in the m/z range from 500 to 4000 was done on a ThermoFisher UHMR system (Thermo Scientific, CA). The sample was loaded into a gold-coated pulled borosilicate and sprayed on the Nanospray Flex Ion Source (Thero Scientific, CA) with spray voltage set as 1.2-1.4 kV. To minimize the gas-phase activation, crucial activation energies and gas pressures were carefully examined and finalized as: in-source trapping (IST): -10V, in-source dissociation (CID): 10V, higher energy in collision cell (HCD): 10V, and trapping gas pressure: 5 (arbitrary unit). The resolution was set to 200,000 (*m/z* 400) and the m/z scan range was from 500 to 20,000 unless otherwise specified.

Direct charge assignment of liraglutide oligomers in high m/z range from 4,000 to 10,000 was done by direct mass technology (DMT) mode embedded in the ThermoFisher UHMR system. The loading of sample and minimization of activation are identical to the abovementioned settings except the trapping gas pressure was set to 0.2 for reduction of collision between neutral ion and gas, an essential condition for individual ion analysis.^28^ Ion filters and charge assignments were done by STORIboard build 1.0.24087.1 (Proteinaceous, IL). The modes of ion transmission and analyzer were set to high and low, respectively.

### ELECTRON-CAPTURE DISSOCIATION (ECD) MASS SPECTROMETRY

Liraglutide solution in the concentration of 1 mg/mL was sprayed on an Agilent 6545 XT Q-ToF system which was configured with a digital quadrupole (DigQ) and an ECD cell as demonstrated in our previous work^24^. Solutions were sprayed on a nanospray source with electrospray voltage set to 1.5 – 1.8 kV. Wide-band isolation on the DigQ was done by altering the duty cycle of 50.0/50.0 to 59.5/40.5 and varying the frequency value based on q-value of 0.59. Electron-capture dissociation was done on an ExDC cell (e-MSion, OR) by using an optimized set of L1-L7 DC bias and filament bias of 11V to maximize fragmentation efficiency, as reported in our previous study.^24^ Generated spectra were deconvoluted by using MASH Native^37^ to convert isotopically resolved peaks into a list of neutral masses. The subsequent peptide mapping was done by using ClipsMS^38^ with a mass tolerance of 20 ppm error.

### MOLECULAR DYNAMICS

Molecular dynamics simulations were performed using GROMACS 2023.3^39^, utilizing the Martini 2.2 force field^40,41^ and the CHARMM36m force field.^42^ The non-standard residue was parametrized by CHARMM-GUI^43^ and formatted using the CGbuilder tool.^44^ Coarse-grained structures were back-mapped to atomic structures using a backward mapping script.^45^ Liraglutide molecules were randomly placed in a periodic cubic box, ensuring a minimum separation of 2 nm. Water beads were mixed with 10% anti-freeze beads as the solvent, and chloride ions were added to neutralize the system. Simulations began with energy minimization, followed by 10 ns of NVT and 10 ns of NPT equilibration. Production simulations were extended to 10 μs. Temperature was controlled by the v-rescale thermostat, and pressure was maintained at 1 bar using the Berendsen barostat for equilibration; Parrinello-Rahman barostat was used for production runs.^46,47^ The 30-mer and 45-mer simulation duplicates were conducted at 300 K and 360 K, generating 8 trajectories in total.

## RESULTS

### Liraglutide Oligomers Characterized by native Mass Spectrometry (nMS)

The size of liraglutide oligomers is highly dependent on solution pH;^6, 9^ however, variations in oligomer sizes have been reported as 6/7/8-mer and 12/13/14-mer in basic and acid conditions, respectively, likely due to the limited resolution of ensemble-based measurements.^6, 8-10^ This limitation can be addressed using the extended mass range and high resolution of the Orbitrap mass spectrometer, which provides unrivaled capabilities for studying large molecular complexes.^14^

To initiate oligomer formation, liraglutide powder was dissolved at pH 6.7 at a high concentration (1 mg/mL) and introduced into the mass spectrometer via a small orifice emitter (diameter∼2 µm), preserving the oligomeric structures. Within 10 min of preparation, both monomeric and oligomeric forms of liraglutide were detected (**Fig. S1**). Initially, oligomers with *n*=2-8 (*m/z* 2000-3500) were more abundant than higher-order oligomers with *n*=13-16 (m/z 3500-5000) (**Fig. 1a**, 10 min). Prolonged incubation resulted in the formation of higher-order oligomers, with a noticeable decrease in the relative abundance of *n*=2-8 oligomers within the m/z range of 2500-3500 (**Fig. 1a**, 30 & 90 min). By 270 min, oligomers with *n*=13-16 became the predominant species. After 24 h of incubation, decreased charge states in *n*=13-16 oligomers were observed, alongside the emergence of a new oligomer (*n*=17) primarily carrying 14 positive charges.

**Figure 1.**
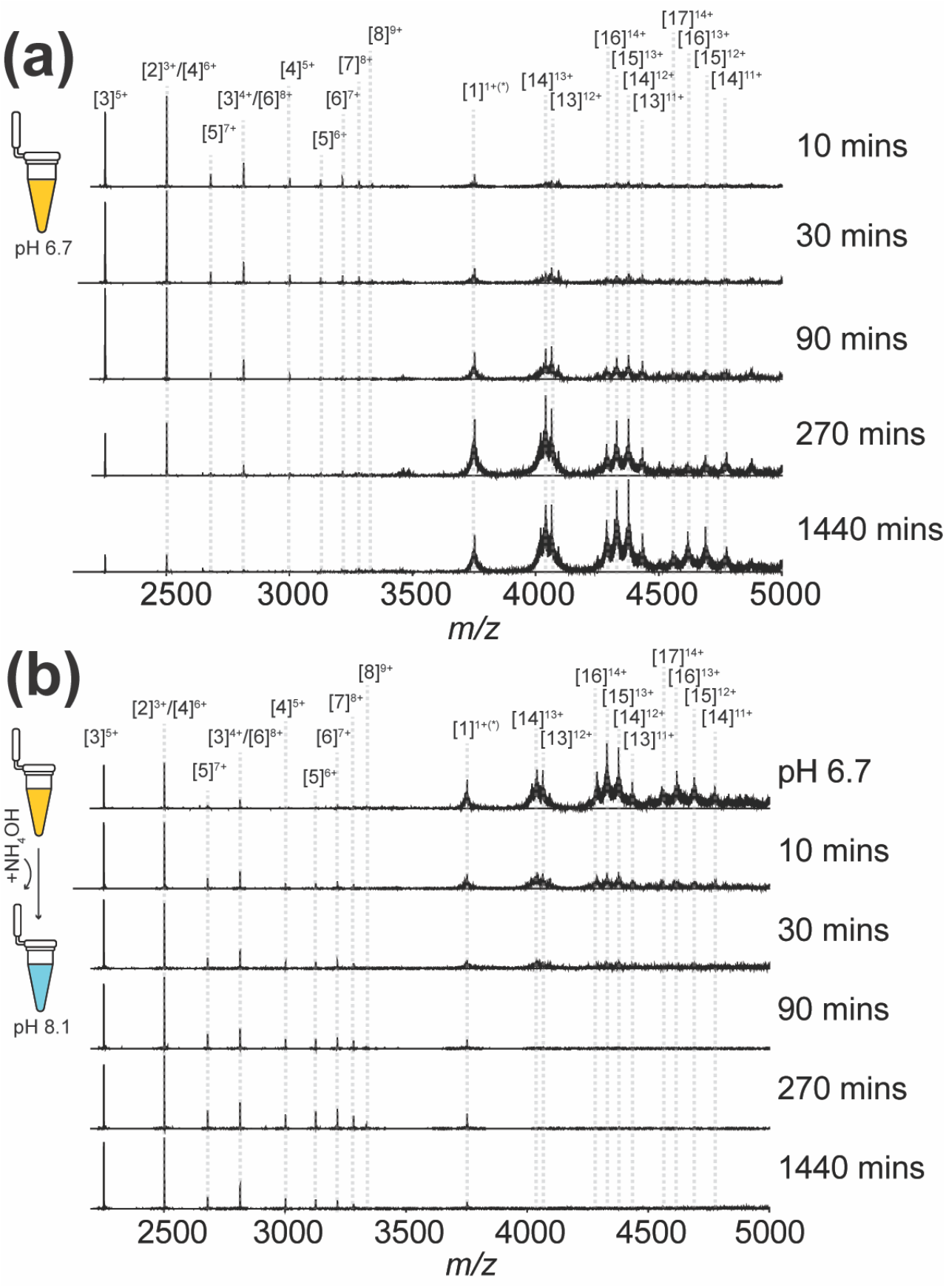
Kinetic monitoring of liraglutide oligomerization in m/z 2000-5000. **(a)** Liraglutide solution incubated and monitored in 20 mM ammonium acetate (AmAc) at pH 6.7. **(b)** Following an 18-hour incubation at pH 6.7, the solution pH was adjusted to 8.1. Oligomers are annotated as [n]^m+^, where n and m represents the oligomeric and charge state, respectively.

To evaluate the reversibility of the oligomerization process, the solution pH was increased from 6.7 to 8.1 following the formation of oligomers in the *n*=13–17 range (**Fig. 1b**). This pH adjustment resulted in an approximately 50% reduction in the relative abundance of these high-order oligomers within 10 min, accompanied by the rapid emergence of oligomers *n*=2-8. These findings indicate that the dissociation of larger oligomers is required for the reformation of smaller oligomers.

The reversible transition between small and large oligomers has been attributed to the protonation state of histidine residues in liraglutide, which modulates the rate of monomer dissociation—a process identified as the rate-limiting step in oligomer transformation.^6^ Beyond the direct interconversion between two oligomeric size classes shown in previous studies, our result further revealed the simultaneous presence of multiple oligomeric states, spanning both the *n*=3–8 and *n*=13–17 ranges. Distinct from proteins, peptides exhibit high structural flexibility and a shallow energy landscape, allowing for a broad distribution of solution conformers.^48, 49^ This structural heterogeneity may therefore underline the capability of liraglutide to form diverse oligomeric states.

### Electron-Capture Dissociation Identified Altered Interfacial Dynamics of Liraglutide Oligomers

We investigated the structural stability of high-order liraglutide oligomers using electron-capture dissociation (ECD). ECD is a fragmentation technique highly sensitive to the structural integrity of proteins and protein complexes and has been widely employed to detect conformational alterations induced by effectors or intermolecular interfaces^24, 50^. To enhance the detection of low-abundance *c* and *z* fragment ions, we employed an advanced digital quadrupole isolation system coupled with an ECD cell, enabling wide-band selection and efficient fragmentation^51^.

To establish a benchmark, we first isolated the 3+ liraglutide ion (*m/z* 1251) for ECD fragmentation (**Fig. 2a**). Unlike conventional collision-induced dissociation (CID), ECD yields a relatively low abundance of fragment ions ∼0.1–0.3% in relative abundance. Notably, despite isolating the 3+ ion, a 2+ ion was also detected at *m/z* 1876, likely resulting from gas-phase charge reduction during electron capture which has been widely noted in ECD experiments.^50^

**Figure 2.**
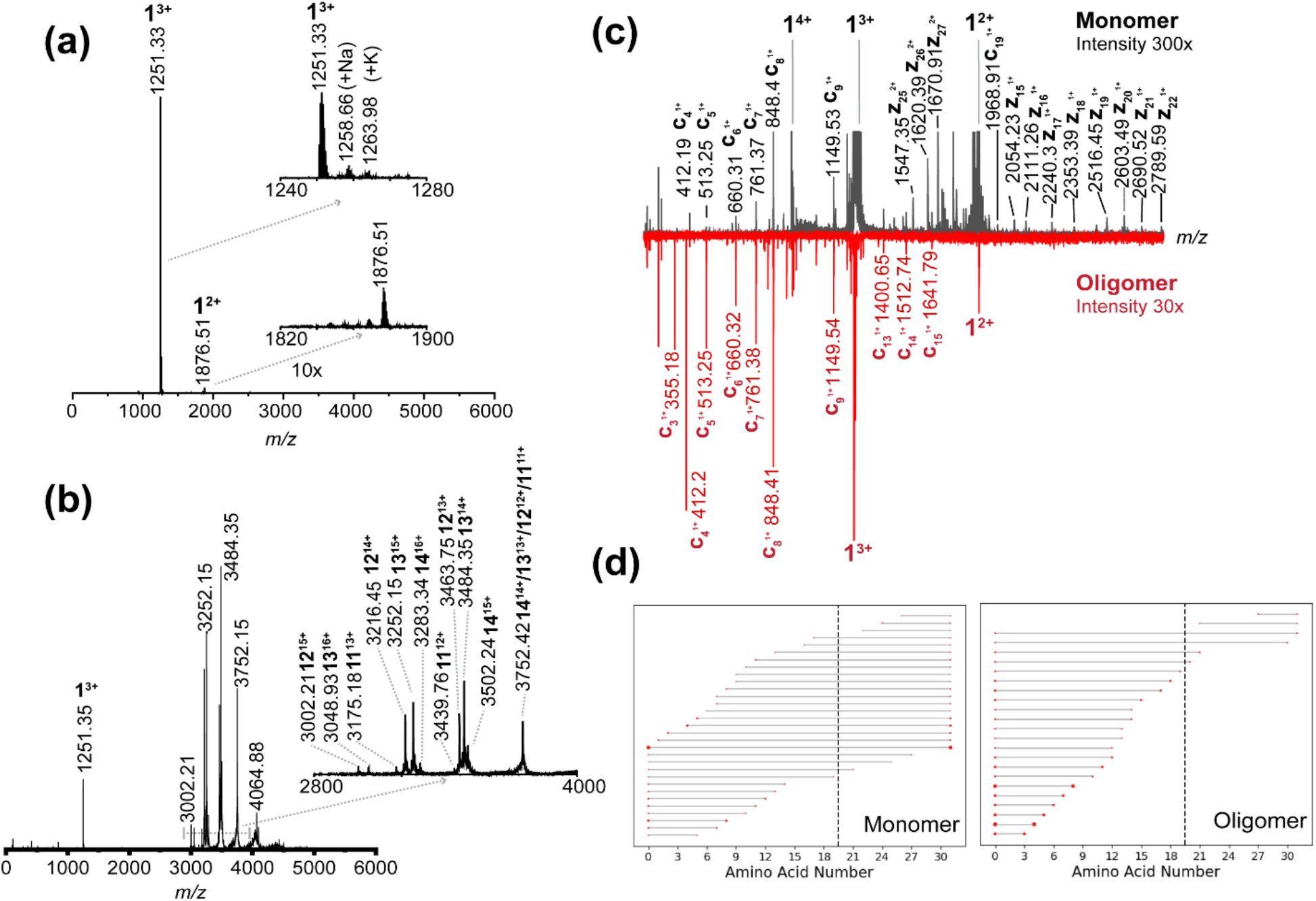
Electron-capture dissociation (ECD) mass spectrometry analysis of liraglutide monomers and oligomers. **(a)** ECD fragmentation of isolated monomeric liraglutide. **(b)** ECD fragmentation of isolated oligomeric (*n*=12-14) liraglutide **(c)** Comparison of monomeric and oligomeric liraglutide fragments in the *m/z* range of 350 to 3000. **(d)** Sequence coverage map showing identified peptide fragments of monomeric and oligomeric liraglutide. The dashed line indicates the lysine residue conjugated to palmitic acid.

To extend this analysis to oligomeric liraglutide, oligomers *n*=11–14 (*m/z* 3000–4000) was isolated and fragmented (**Fig. 2b**). Notably, despite maintaining identical solution conditions (pH 6.7, 20 mM ammonium acetate), the oligomeric distributions observed in the Q-ToF system differed slightly from those detected in the Q-Orbitrap system, with fewer oligomers preserved in the Q-ToF analysis. This discrepancy likely arises from suboptimal ion transmission in the current Q-ToF configuration. Although such variations highlight potential differences arising from activation parameters or instrument design, both mass spectrometers consistently revealed the heterogeneous oligomeric states that were previously unresolved by conventional analytical methods. These findings underscore the power of mass spectrometry in resolving highly complex oligomeric assemblies within a single analysis.

Fragmentation analysis revealed that both monomeric and oligomeric liraglutide primarily generated singly or doubly charged c and z ions distributed across the *m/z* range of 400–2700 (**Fig. 2c**). Notably, fragmentation efficiency was higher in oligomeric liraglutide due to the higher charge states facilitating the electron-capture process. However, in the *m/z* 2000–2700 range, a reduction in *z* ions was observed for oligomeric liraglutide (**Fig. 2c**, bottom panel).

To correlate these fragmentation patterns with structural features, we mapped the product ions onto a sequence coverage map of liraglutide (**Fig. 2d**). The coverage map clearly demonstrates that oligomeric liraglutide yielded fewer *c-, z-*fragments compared to monomeric liraglutide. Additionally, the amount of *c-* and *z-*ions with palmitic acid conjugates were less evident in oligomeric liraglutide, suggesting the conjugation limits fragmentation when oligomer formed. Beyond the conjugation site, an unexpected reduction in C-terminal *z* ions was observed, indicating restricted dynamics in the C-terminal domain upon oligomerization (**Fig. 2d**). These results suggest that oligomerization is primarily driven by hydrophobic interactions alongside restricted dynamics of C-terminal residues, yielding stable self-assembled oligomeric structures.

### Newly Discovered Higher Oligomeric States of Liraglutide Observed by Direct Mass Technology^TM^ Measurement

In addition to the previously reported soluble oligomers, we detected low-abundance species in the *m/z* range of 5,000–10,000 after an 18-hour incubation in 20 mM ammonium acetate (pH 6.7). However, the overlapping signals within this range produced low-resolution spectra with convoluted peak distributions, precluding accurate charge state determination^52^ using conventional charge state distribution analysis (**Fig. 3a**). This limitation was overcome by using Direct Mass Technology (DMT), a method in which the accumulation of ion signal intensity over analysis time is proportional to the ion charge,^27, 28, 53^ enabling charge assignment by analyzing the slope of summed signal intensity versus analysis time.

**Figure 3.**
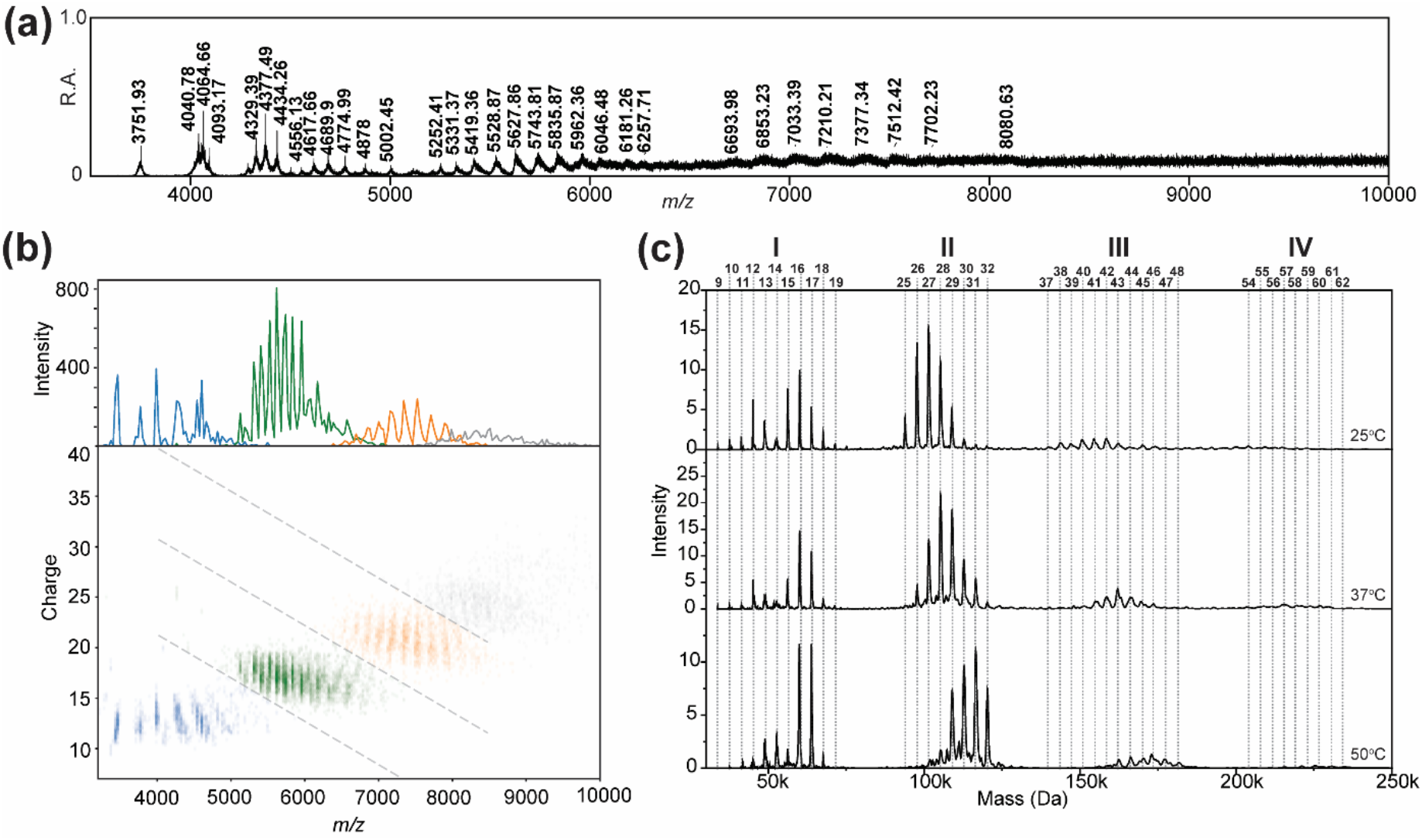
Liraglutide oligomers in high mass range. **(a)** Mass spectra acquired in conventional MS mode **(b)** Two-dimensional (2D) heatmap analysis of data acquired in DMT mode. The charge state of induvial ion was assigned using STORI analysis. Clusters are color-coded based on the distinct regions (I-IV) observed in (c). **(c)** Direct mass spectra showing four distinct regions (I-IV) at different incubation temperatures. Oligomeric states are indicated on the top of the spectra.

Using DMT, we determined that the charge states of these high *m/z* species primarily ranged from 8+ to 75+ (**Fig. 3b**). Crucially, we found that optimizing charge assignment parameters, such as narrowing the bin size and reducing the minimum number of ions required for analysis, was essential to retain sufficientions for subsequent charge assignments. This was not achievable using the default STORI charge assignment parameters (**Fig. S2**). Visualization of *m/z* (x-axis) versus charge (y-axis) revealed four distinct charge-state clusters, which were color-coded in the raw spectra for validation (**Fig. 3b**). Mapping these color assignments back to their respective *m/z* domains clearly delineated overlapping signal regions, highlighting the complexity of charge distribution in this *m/z* range (**Fig. 3b**, orange and gray). This charge assignment enabled mass determination of the newly identified oligomers, which ranged from ∼100 kDa to 250 kDa, corresponding to oligomeric states of *n*=25–62 (**Fig. 3c**).

Interestingly, rather than forming a continuous distribution of oligomeric states from *n*=12 to *n*=62, these high molecular weight (HMW) oligomers segregated into four distinct clusters: *n*=9–19 (Region I), *n*=25– 32 (Region II), *n*=37–48 (Region III), and *n*=54–62 (Region IV) (**Fig. 3c**). To investigate the influence of solution conditions on the distribution of these HMW oligomers, we evaluated the effects of pH and temperature. While HMW oligomers were undetectable at elevated pH (**Fig. S3**), increasing the temperature at pH 6.7 promoted oligomer growth across all four regions (**Fig. 3c**). In Region I, a bimodal distribution emerged, with two dominant states (*n*=12 and *n*=16) at 25 °C. At higher temperatures (37 °C and 50 °C), the distribution became more uniform, with *n*=16 as the dominant species. Notably, this bimodal distribution was exclusive to Region I, whereas in the remaining three regions, elevated temperatures led to a general shift toward higher oligomeric states. The distinct segregation of oligomeric clusters and the unusual bimodal distribution in Region I prompted further investigation via molecular dynamics simulations to elucidate the underlying mechanisms driving these structural phenomena.

### Molecular Dynamic Simulation Proposed a Plausible Oligomerization Mechanism Coherent with DMT Observation

To elucidate the formation of high-mass oligomers observed in our experiments, we performed a 10-µs coarse-grained molecular dynamics (MD) simulation with 30 liraglutide monomers. The monomers were initially randomized within a confined 26.22 nm^3^ box with explicit water molecules (liraglutide concentration∼3 mM) and assigned charges based on their propensity to carry charges at pH 6.7.

As an amphiphilic peptide^54^, liraglutide undergoes self-assembly driven by a balance between hydrophobic and hydrophilic interactions. To determine their individual contributions to oligomerization, predefined hydrophobic and hydrophilic residues were color-mapped in red (hydrophobic) and blue (hydrophilic) (**Fig. 4a**). At 0 µs, each monomer exhibited a distinct segregation of hydrophobic and hydrophilic regions, primarily attributed to the hydrophobicity of D6M and the hydrophilicity of backbone residues (**Fig. 4a**, 0 µs). As the simulation progressed, liraglutide monomers associated into small, transient oligomers (*n*=5–8) via hydrophobic interactions (**Fig. 4a**, 0.4 µs). These oligomers evolved into larger oligomers through the fusion of two oligomers. Interestingly, unlike the assembly process of small oligomer formation, the generation of large oligomers (*n*=12-15) required structural rearrangement of the micelle-like core, as evidenced by the presence of dispersed hydrophobic cores which then converged to defined hydrophobic cores (**Fig. 4a**, 3 µs). By the end of the simulation (10 µs), two distinct clusters had formed, exhibiting a clear hydrophilic interface and separated dense hydrophobic centers. Collectively, the growth of large oligomers ceased once a critical size was reached, at which point *n*=12-15 oligomers acted as building blocks for higher order assemblies driven by hydrophilic interactions. MD simulations provide a mechanistic explanation for the discontinued oligomeric states observed beyond *n*=19 and the origins of the higher-order oligomers (*n*=25-32, 37-48, and 54-62) identified by our direct mass measurements.

**Figure 4.**
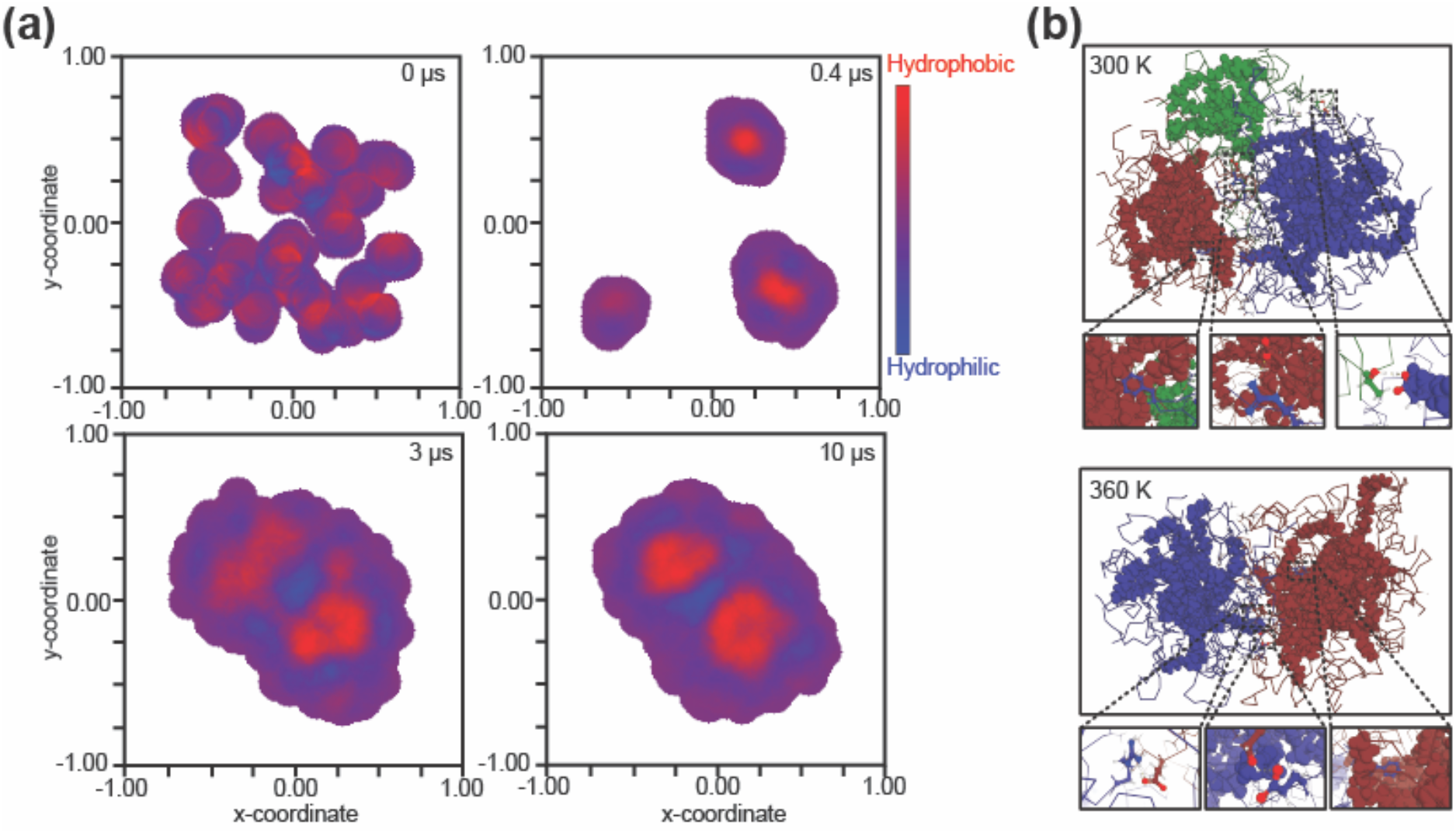
Representative frames and structures from molecular dynamics simulations of liraglutide self-assembly. **(a)** Snapshots at 0, 0.4 µs, 3 µs, and 10 µs show the assembly process of 30 liraglutide monomers. Hydrophobic and hydrophilic regions are labeled in red and blue, respectively. **(b)** Structural comparison of the 30-mer at 300 K and 360 K (10 µs). Peptide backbones (ribbon) and conjugated palmitic acid motifs (spheres) are shown. The number of clusters, determined by the number of dense hydrophobic cores shown in (a), is visualized using green, red, and blue: three clusters at 300 K and two at 360 K. Representative residue pair interactions are shown in the insets.

A detailed contribution of individual residue was analyzed to investigate the molecular forces stabilizing these assemblies (**Fig. 4b**). Three types of interaction pairs were classified: hydrophobic, hydrophilic, and hybrid. For example, van der Waals interactions between phenylalanine and the palmitic acid tail were categorized as hydrophobic pair interactions (**Fig. 4b**, left panel). Analogously, the short distance (<4.5Å) between histidine and glutamic acid was recognized and defined as hydrophilic pair interactions. The final category, hybrid pairs, accounted for interactions between polar and nonpolar residues, which were predominantly stabilized by hydrogen bonding and polarizable side-chain interactions. Unlike the process observed for small (*n*=2-8) and large oligomers (*n*=12-18), in these high molecular weight oligomers, analysis of interfacial interactions revealed a strong contribution from 32-61% hydrophilic-hydrophilic and 20-55% hybrid interactions (**Table S1**). Residue analysis alongside the absence of hydrophobic core rearrangement during the assembly of high-order oligomers suggested that electrostatic and hydrogen bond networks on the surface are the primary driving force in their association.

As the simulation corroborated our DMT findings, we extended this approach to investigate the effects of temperature. In addition to the baseline simulation at 300 K, we conducted additional simulations at 360 K while maintaining all other parameters constant. Simulation at each temperature was performed in duplicate to ensure the reliability of our observations. By the end of the simulations, all 30 liraglutide monomers had assembled into a large 30-mer, with minor variations in intermediate oligomeric states (**Fig. S4**). Notably, at 300 K, two- and three-distinct clusters formed, with oligomeric sizes of 8-22 and 5-10-15 (where x-y-z denotes the number of liraglutide molecules in each cluster). At 360 K, the simulations yielded two clusters with oligomeric sizes of 12-18 and 13-17. (**Fig. 4b**). Extended study using 45 liraglutide monomers showed the clusters of 9-10-11-15, 5-6-14-20 and 8-9-10-18, 10-17-18 at 300 K and 360 K, respectively. (**Fig. S5**) These results highlighted that the increase of temperature yielded a more uniform distribution of the large oligomers as building blocks. Detailed examination of the course of simulation showed that elevated temperatures promote the growth of *n*=9–18 oligomers into larger assemblies by increasing peptide backbone flexibility and facilitating structural rearrangement. Additionally, high temperature promoted an increased frequency of the formation and disruption of interfacial contacts, enabling oligomers with suboptimal interactions (high-energy states) to dissociate and reassociate into more stable oligomeric form. The resulting observation provided explanatory evidence of our DMT observation where a more uniform distribution and larger oligomers were observed at 37 ^o^C and 50 ^o^C compared to 25 ^o^C in Region I (*n*=9-18). Subsequently, the increased size of these *n*=9-18 building block oligomers at elevated temperatures contributes to the overall growth of high-order oligomers (*n*=25-32, 37-48, and *n*=54-62) through the previously observed hydrophilic interactions, resulting in the observed increase oligomeric states in Region II-IV detected by DMT.

## DISCUSSION AND CONCLUSION

Conjugating therapeutic peptides to generate amphiphilic properties has promoted successful drug delivery, enhanced pharmacokinetics, and increased stability in the human circulatory system.^55-57^ However, the propensity for oligomerization can induce insoluble aggregates, adversely affecting both the manufacturing process and biomedical function. The formation of insoluble species has been identified by transmission electron microscopy forming micrometer-scale fibrillar or amorphous aggregates.^9^ While identification of these oligomers is crucial since they are potential precursors or inhibitors of the final precipitated insoluble products^58^, stress studies failed to correlate the morphology of insoluble aggregates to the composition of pre-fibrillation oligomers.^9^ Native mass spectrometry (nMS) offers a highly specific approach analyzing hidden soluble oligomers and deciphering the underlying mechanisms of their formation. Advancements in fragmentation and detection in native mass spectrometry broadens the scope of oligomer analysis, including identification of the oligomers’ interfacial dynamics and the exploration of previously hidden species.

Native mass spectrometry can resolve highly heterogeneous systems to reveal detailed insights across a wide range of mass domains. At *m/z* ranges less than 3000, corresponding to oligomeric states *n*=2 to 8 of liraglutide, masses were unambiguously determined by isotopically resolved signals offered by high-resolution Orbitrap mass spectrometry. In *m/z* 3000 to 5000, oligomers identities were resolved based on their charge state distribution (CSD). However, high heterogeneity and low abundance of species posed challenges to confident determination of species beyond *m/z* 5000. This challenge was addressed by extending the analysis from conventional MS to direct mass technology (DMT) mode, enabling direct assignment of charges to individual ions followed by mass determination. This unambiguous assignment of oligomers allowed us to monitor the assembly trajectory along the course of incubation. On top of the oligomers from *n*=2 to *n*=17, novel liraglutide oligomers *n*=25 to 62 shown in DMT directed a new oligomerization mechanism distinct from that involving hydrophobic conjugates.

Combining electron-capture dissociation (ECD) and molecular dynamics simulations (MDS) provided mechanistic insights into the oligomerization pathway following the formation of oligomers with *n*=12 to 18. ECD was used to probe the structural dynamics of these oligomers, while MDS provided a detailed view of their interactions and assembly. The decreased release of z ions observed after the formation of 12 to 14-mers was interpreted as an indication of restricted C-terminal residue dynamics in larger oligomers. Based on ECD and MDS results, we propose two essential processes for the evolution of *n*=12-18 oligomers into higher-order (*n*=25-62) oligomers: (i) restriction of C-terminal residues stabilizes the structures of *n*=12-18 oligomers and (ii) subsequent association with hydrophilic residues leads to the formation of segregated clusters, as detected by direct mass spectrometry. The agreement between our experimental and simulation results promoted us to conduct a comprehensive examination of the liraglutide oligomerization process, with a focus on the conformational and energy landscapes adopted during oligomerization. These findings are presented in a parallelly submitted manuscript.

Identifying the intermediate oligomers is challenging and requires advanced analytical platforms. Distinct from folded protein in the biological system, the inherent flexibility of peptide backbones allows them to adopt a wide range of conformers. The conjugation of hydrophobic moiety further diversifies the conformational landscape of peptide therapeutics. Uncertainty regarding conformational distribution can cascade the off-pathway oligomerization and eventually insoluble aggregates. Although dual oligomerization pathways are commonly described in controlling the morphology of nanomaterials formed from amphiphilic building blocks,^59-61^ studies explicitly noting this phenomenon in the context of therapeutic conjugates are limited. While current studies extensively focus on oligomers formed via hydrophobicity imparted by conjugated lipid tails, we identified higher-order oligomers formed through a combination of hydrophobic and hydrophilic interactions. The presence of these higher-order oligomers may be more indicative of fibrillation observed by microscopic analysis.^62, 63^ Parallel to other analytical techniques, we demonstrated a promising array of mass spectrometric techniques to identify uncommon oligomers in solution, supported by simulation data. We hope this will provide a novel framework for the discovery of unexpected high-order oligomers, which may risk aggregation or reduced efficacy, in the field of drug discovery, design, and development.

## SUPPORTING INFORMATION

Full range mass spectra, optimization of direct mass charge assignment parameters, analysis of high-order oligomers at high pH condition, molecular dynamics process of high-order oligomers, summary of structures of liraglutide oligomers analyzed by MD

## ACKNOWLEDGMENT

The authors acknowledge funding and material support from Boehringer Ingelheim Pharmaceuticals, Inc., National Institutes of Health (R35GM143047 to X. Y. and RM1GM149374 to D.H.R.), and the Robert A. Welch Foundation (Grant A-2089 to X.Y. and Grant A-2162 to D.H.R.).

## TOC

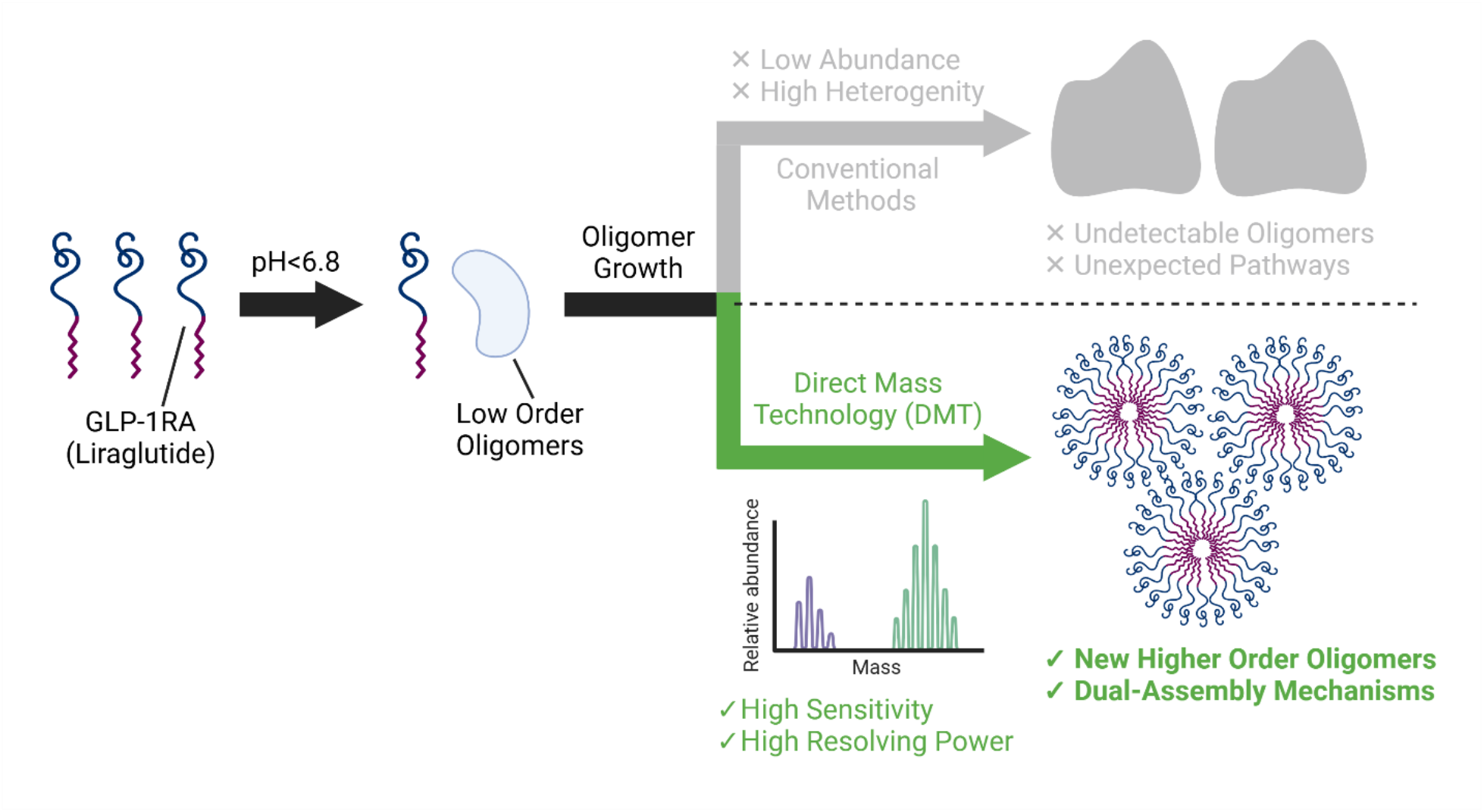

