## Supporting information for "Unveiling the Hidden: Dissecting Liraglutide Oligomerization Dual Pathways via Direct Mass Technology, Electron-Capture Dissociation, and Molecular Dynamics"

Table of Contents

| | Page |
|---|---|
| S1. Full Mass Range Analysis of Liraglutide Oligomers | |
| Figure S1 | S2 |
| Table S1 | S3 |
| S2. Optimization of Charge Assignment Parameters for Liraglutide Analysis | |
| Table S2 | S4 |
| Figure S2 | S4 |
| S3. Analysis of High-Order Oligomers at pH 6.7 and 8.1 | |
| Figure S3 | S5 |
| S4. Molecular Dynamics Simulation of 30 Liraglutide Monomer Assembly | |
| Figure S4 | S6 |
| S5. Interaction Network of High-Order Oligomers from Molecular Dynamics Simulations | |
| Table S4 | S7 |
| Table S5 | S8 |

### S1. Full Mass Range Analysis of Liraglutide Oligomers

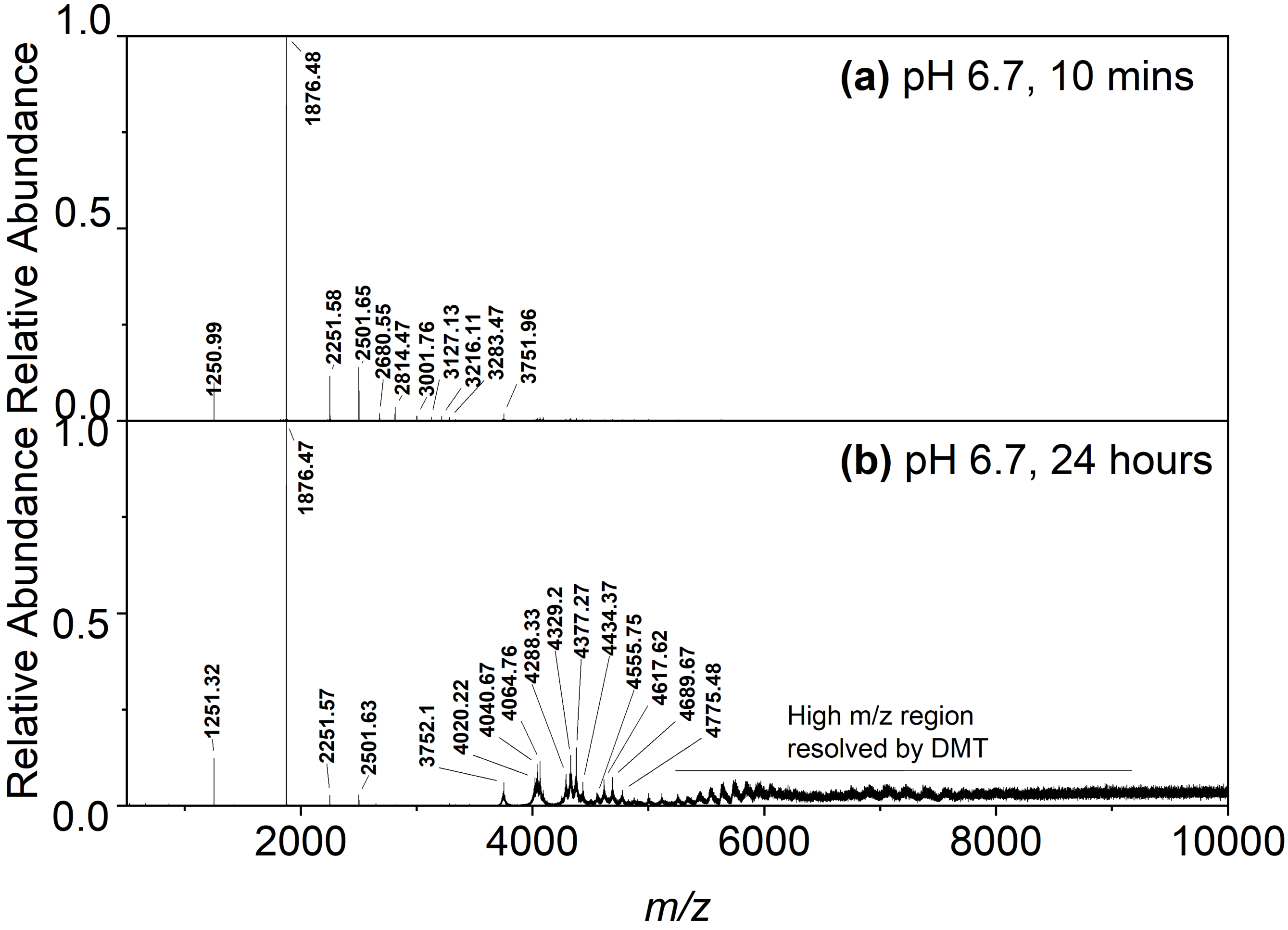


**Figure S1.** Full mass range (*m/z* 500-10000) of mass spectra of liraglutide in the solution at pH 6.7 after (a) 10 min and (b) 24 h. Liraglutide masses were determined from isotopically resolved peaks or charge state distribution and summarized in **Table S1**.

| m/z | charge | mass (Da) | oligomeric states (n) | Mass Determination |
|---|---|---|---|---|
| 1250.99 | 3 | 3749.958 | 1 | Isotopic distribution |
| 1876.48 | 2 | 3750.952 | 1 | Isotopic distribution |
| 2251.58 | 5 | 11252.88 | 3 | Isotopic distribution |
| 2501.65 | 3 | 7501.938 | 2 | Isotopic distribution |
| 2680.55 | 7 | 18756.822 | 5 | Isotopic distribution |
| 2814.47 | 8 | 22507.728 | 6 | Isotopic distribution |
| 3001.76 | 5 | 15003.78 | 4 | Isotopic distribution |
| 3127.13 | 6 | 18756.756 | 5 | Isotopic distribution |
| 3216.11 | 7 | 22505.742 | 6 | Isotopic distribution |
| 3283.47 | 8 | 26259.728 | 7 | Isotopic distribution |
| 3751.96 | 1/14 | 3750.956/ 52513.384 | 1/14 | Charge state distribution |
| 4020.22 | 14 | 56269.024 | 15 | Charge state distribution |
| 4040.67 | 13 | 52515.658 | 14 | Charge state distribution |
| 4064.76 | 12 | 48765.072 | 13 | Charge state distribution |
| 4288.33 | 14 | 60022.564 | 16 | Charge state distribution |
| 4329.2 | 13 | 56266.548 | 15 | Charge state distribution |
| 4377.27 | 12 | 52515.192 | 14 | Charge state distribution |
| 4434.37 | 11 | 48767.026 | 13 | Charge state distribution |
| 4555.75 | 14 | 63766.444 | 17 | Charge state distribution |
| 4617.62 | 13 | 60016.008 | 16 | Charge state distribution |
| 4689.67 | 12 | 56263.992 | 15 | Charge state distribution |
| 4775.48 | 11 | 52519.236 | 14 | Charge state distribution |

**Table S1.** Masses, charge, and oligomeric states of liraglutide oligomers determined by isotopically resolved peaks or charge state distribution.

### S2. Optimization of Charge Assignment Parameters for Liraglutide Analysis

| | (a) Default | (b) Narrow window |
|---|---|---|
| Parameter | | |
| Bin size (ppm) | 1.5 | 1 |
| Minimum Ions in Bin | 2 | 1 |
| Numbers of Charge Neighbors | 2 | 2 |
| Number of Isotope Neighbors | 10 | 10 |
| Result | | |
| Assigned ion counts | 389 | 11,290 |
| Assigned charge range | 11-19 | 8-75 |

**Table S2.** Optimization of charge assignment parameters in the STORI voting v3 charge assigner. (a) Default parameters with a wide bin size resulted in excessive ion filtering. (b) Narrowing the bin size reduced ion filtering, while adjusting the minimum number of ions per bin compensated for the reduced number of ions in each bin. A total of 2,702,223 ions were analyzed and filtered using both STORI processor and charge assignment parameters to minimize false positive identifications.

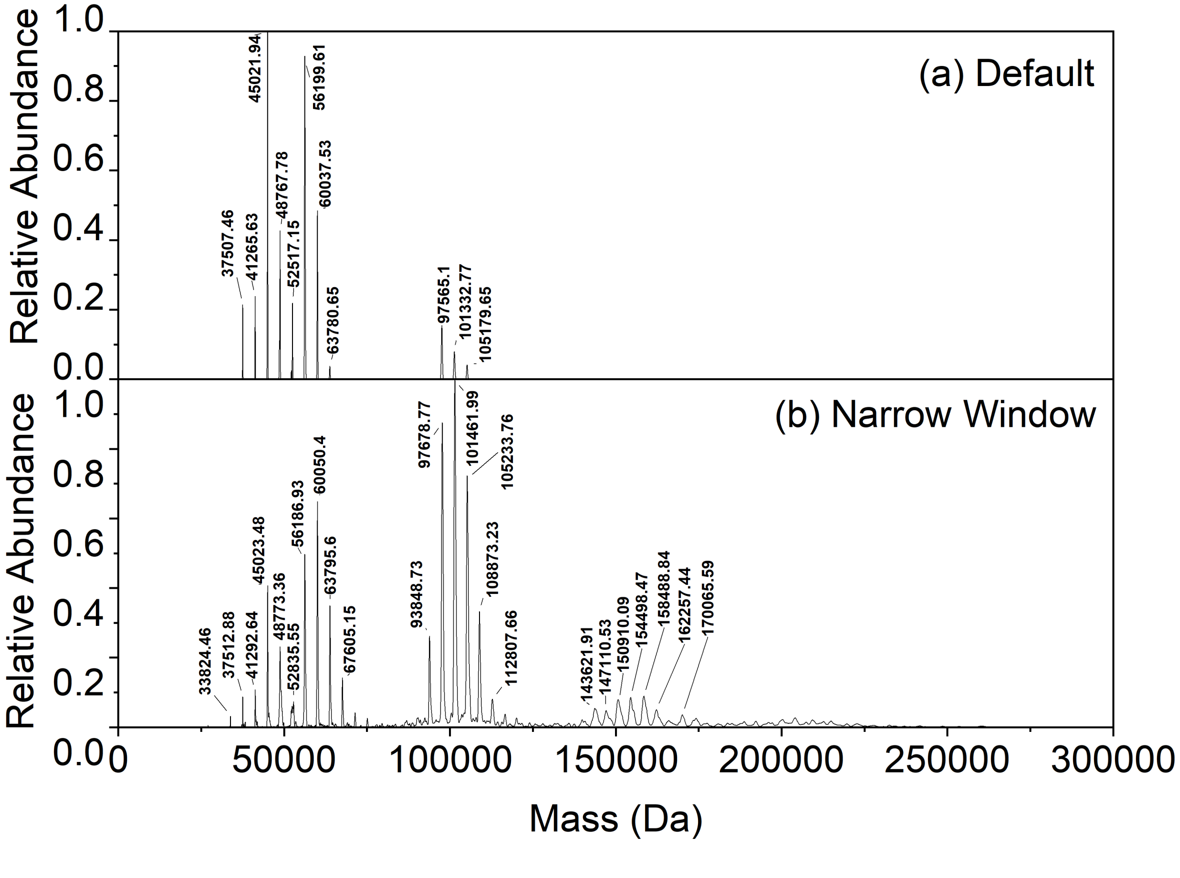


**Figure S2.** Direct mass spectra of liraglutide at pH 6.7 and 25 ºC resulting from the charge assignment using (a) default and (b) narrow bin size windows

### S3. Analysis of High-Order Oligomers at pH 6.7 and 8.1

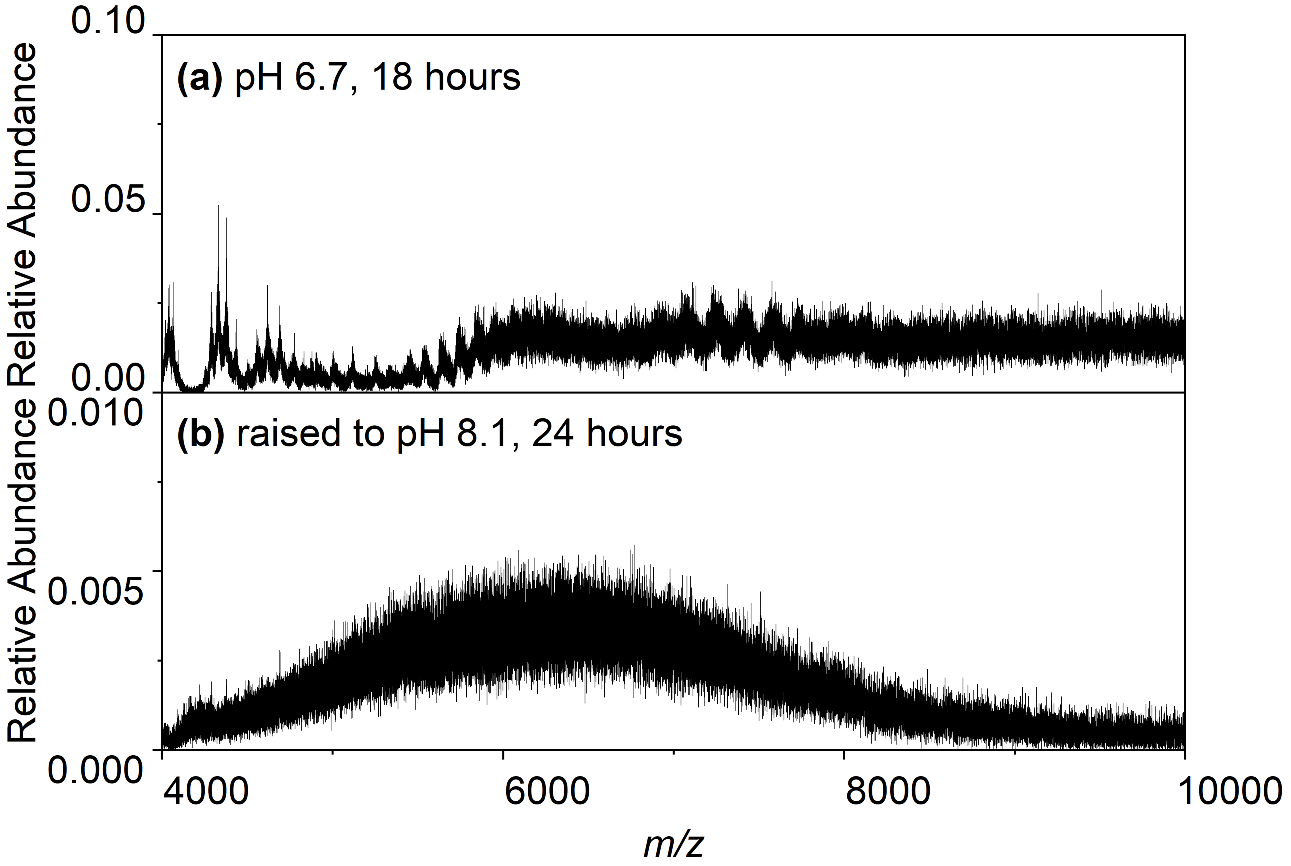


**Figure S3.** Zoomed mass spectra (m/z 4,000-10,000) of liraglutide in solution at (a) pH 6.7, incubated 18 h (initial condition) and (b) pH 8.1 after 24 h of incubation.

### S4. Molecular Dynamics Simulation of 30 Liraglutide Monomer Assembly

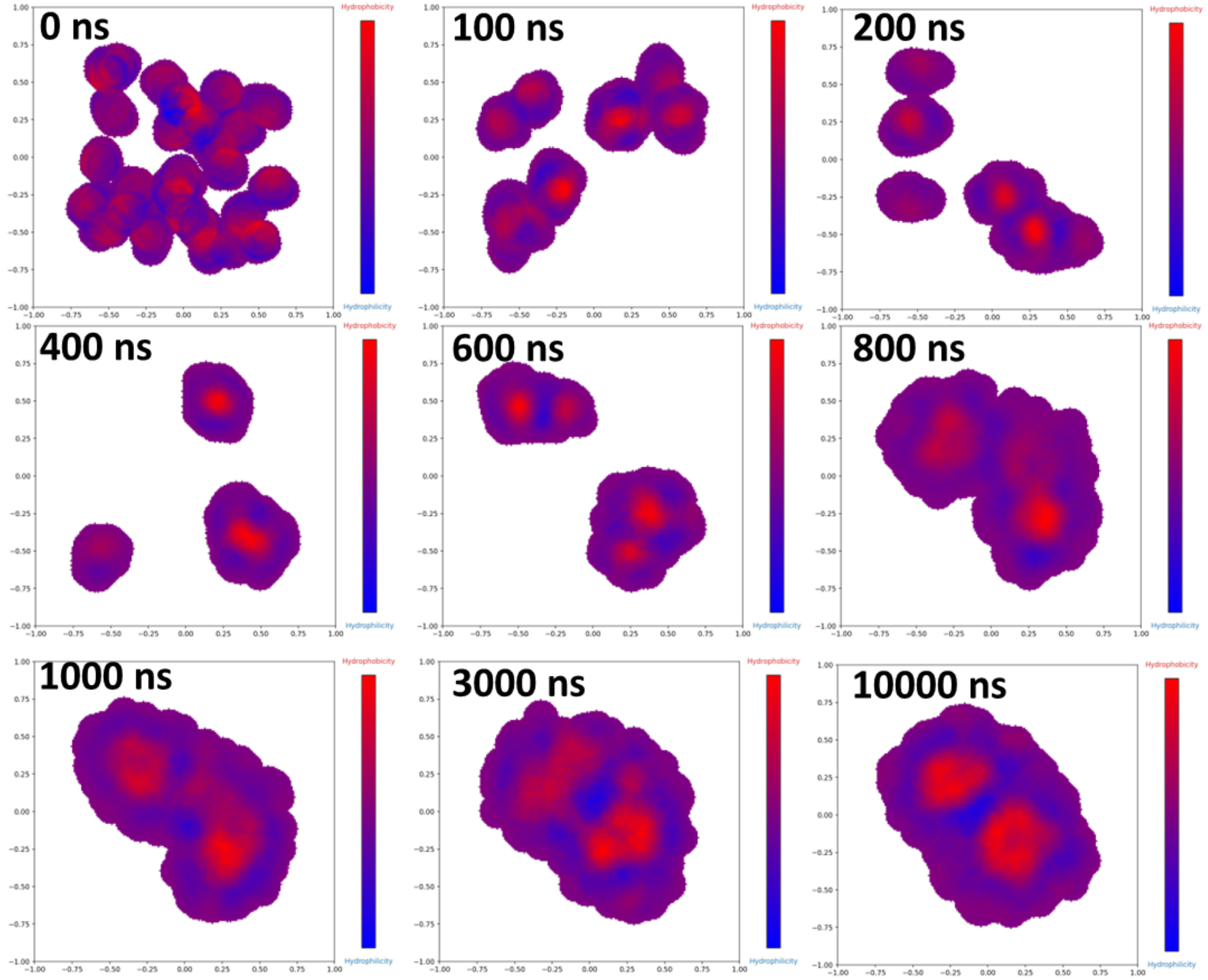


**Figure S4.** Molecular dynamic simulation of 30 liraglutide molecules assembling at 300 K. Hydrophobic and hydrophilic regions are shown in red and blue, respectively.

### S5. Interaction Network of High-Order Oligomers from Molecular Dynamics Simulations

**Table S4.** Summary of end-point structure from eight simulation results. Subunit composition indicates the number of liraglutide monomers in each subunit. Contact contributions were calculated from residue statistics presented in **Table S5**.

| Condition | Structure | Hydrophobic Contacts Contribution (%) | Hydrophilic Contacts Contribution (%) | Hybrid Contacts Contribution (%) | Subunit Composition (w-x-y-z) |
|---|---|---|---|---|---|
| 30mer-300K-Rep1 | 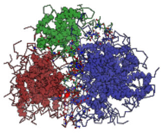 | 22 | 32 | 46 | 5-10-15 |
| 30mer-300K-Rep2 | 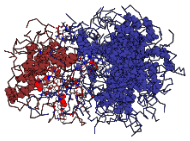 | 22 | 35 | 43 | 8-22 |
| 30mer-360K-Rep1 | 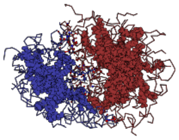 | 6 | 61 | 33 | 12-18 |
| 30mer-360K-Rep2 | 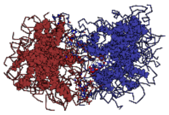 | 27 | 53 | 20 | 13-17 |
| 45mer-300K-Rep1 | 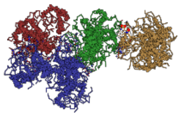 | 19 | 36 | 45 | 9-10-11-15 |
| 45mer-300K-Rep2 | 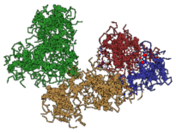 | 16 | 55 | 29 | 5-6-14-20 |
| 45mer-360K-Rep1 | 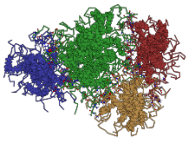 | 10 | 35 | 55 | 8-9-10-18 |
| 45mer-360K-Rep2 | 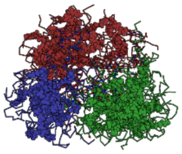 | 12 | 47 | 41 | 10-17-18 |

**Table S5.** Interfacial residue pairs in **(a)** 30-mer, 300 K, replicate 1

| Entry | Cluster Pair | Chain 1 | Res1# | Res1 Name | Chain 2 | Res2# | Res2 Name | Distance (Å) |
|---|---|---|---|---|---|---|---|---|
| 1 | 1-2 | J | 10 | VAL | F | 24 | ALA | 4.11 |
| 2 | 1-2 | J | 11 | SER | F | 30 | ARG | 4.41 |
| 3 | 1-2 | K | 12 | SER | d | 5 | THR | 4.39 |
| 4 | 1-2 | T | 24 | ALA | D | 11 | SER | 4.46 |
| 5 | 1-2 | Y | 10 | VAL | N | 4 | GLY | 3.94 |
| 6 | 1-2 | Y | 11 | SER | N | 4 | GLY | 3.87 |
| 7 | 1-2 | Y | 15 | GLU | N | 1 | HIS | 4.47 |
| 8 | 1-2 | Y | 18 | ALA | D | 5 | THR | 3.58 |
| 9 | 1-2 | Y | 19 | ALA | D | 18 | ALA | 3.97 |
| 10 | 1-2 | Y | 24 | ALA | N | 5 | THR | 4.49 |
| 11 | 1-2 | Y | 24 | ALA | N | 10 | VAL | 3.98 |
| 12 | 1-2 | a | 4 | GLY | F | 30 | ARG | 4.50 |
| 13 | 1-2 | a | 8 | SER | F | 27 | VAL | 4.44 |
| 14 | 1-2 | a | 9 | ASP | d | 7 | THR | 3.86 |
| 15 | 1-2 | b | 24 | ALA | D | 10 | VAL | 4.40 |
| 16 | 1-2 | b | 29 | GLY | D | 8 | SER | 3.75 |
| 17 | 1-3 | J | 28 | ARG | X | 3 | GLU | 4.28 |
| 18 | 1-3 | J | 29 | GLY | X | 5 | THR | 4.41 |
| 19 | 1-3 | L | 17 | GLN | M | 14 | LEU | 4.47 |
| 20 | 1-3 | L | 19 | ALA | M | 11 | SER | 4.32 |
| 21 | 1-3 | R | 2 | ALA | P | 3 | GLU | 4.34 |
| 22 | 1-3 | R | 2 | ALA | P | 4 | GLY | 3.81 |
| 23 | 1-3 | T | 10 | VAL | W | 7 | THR | 4.03 |
| 24 | 1-3 | T | 18 | ALA | M | 19 | ALA | 4.20 |
| 25 | 1-3 | T | 22 | PHE | M | 20 | D6M | 1.95 |
| 26 | 1-3 | T | 30 | ARG | W | 4 | GLY | 3.97 |
| 27 | 1-3 | b | 7 | THR | P | 2 | ALA | 3.82 |
| 28 | 1-3 | c | 9 | ASP | M | 2 | ALA | 4.19 |
| 29 | 1-3 | c | 11 | SER | M | 2 | ALA | 4.04 |

| | | | | | | | | |
|---|---|---|---|---|---|---|---|---|
| 30 | 2-3 | D | 9 | ASP | M | 1 | HIS | 4.05 |
| 31 | 2-3 | D | 9 | ASP | M | 10 | VAL | 3.89 |
| 32 | 2-3 | D | 14 | LEU | P | 10 | VAL | 4.49 |
| 33 | 2-3 | D | 28 | ARG | E | 19 | ALA | 4.27 |
| 34 | 2-3 | N | 2 | ALA | X | 7 | THR | 4.17 |
| 35 | 2-3 | N | 16 | GLY | U | 3 | GLU | 3.78 |
| 36 | 2-3 | N | 19 | ALA | O | 19 | ALA | 3.75 |
| 37 | 2-3 | V | 9 | ASP | E | 24 | ALA | 3.87 |

**(b)** 30-mer, 300K, replicate 2

| Entry | Cluster Pair | Chain 1 | Res1# | Res1 Name | Chain 2 | Res2# | Res2 Name | Distance (Å) |
|---|---|---|---|---|---|---|---|---|
| 1 | 1-2 | B | 5 | THR | a | 4 | GLY | 3.94 |
| 2 | 1-2 | G | 2 | ALA | J | 3 | GLU | 4.37 |
| 3 | 1-2 | G | 2 | ALA | J | 5 | THR | 3.65 |
| 4 | 1-2 | G | 3 | GLU | J | 2 | ALA | 3.89 |
| 5 | 1-2 | G | 8 | SER | K | 2 | ALA | 3.49 |
| 6 | 1-2 | N | 7 | THR | c | 21 | GLU | 4.39 |
| 7 | 1-2 | N | 8 | SER | c | 24 | ALA | 3.89 |
| 8 | 1-2 | N | 10 | VAL | c | 24 | ALA | 4.19 |
| 9 | 1-2 | N | 11 | SER | c | 29 | GLY | 3.59 |
| 10 | 1-2 | N | 20 | D6M | R | 13 | TYR | 4.26 |
| 11 | 1-2 | Q | 27 | VAL | c | 24 | ALA | 4.23 |
| 12 | 1-2 | S | 8 | SER | P | 20 | D6M | 4.49 |
| 13 | 1-2 | S | 13 | TYR | V | 20 | D6M | 3.75 |
| 14 | 1-2 | S | 15 | GLU | R | 2 | ALA | 3.89 |
| 15 | 1-2 | S | 16 | GLY | R | 2 | ALA | 3.70 |
| 16 | 1-2 | S | 16 | GLY | R | 5 | THR | 3.58 |
| 17 | 1-2 | S | 20 | D6M | R | 10 | VAL | 1.66 |
| 18 | 1-2 | T | 4 | GLY | J | 1 | HIS | 4.32 |
| 19 | 1-2 | T | 12 | SER | V | 30 | ARG | 4.12 |

| | | | | | | | | |
|---|---|---|---|---|---|---|---|---|
| 20 | 1-2 | U | 7 | THR | R | 3 | GLU | 4.38 |
| 21 | 1-2 | U | 7 | THR | R | 4 | GLY | 4.24 |
| 22 | 1-2 | Z | 4 | GLY | P | 19 | ALA | 4.01 |
| 23 | 1-2 | Z | 11 | SER | P | 24 | ALA | 3.88 |

**(c)** 30-mer, 360K, replicate 1

| Entry | Cluster Pair | Chain 1 | Res1# | Res1 Name | Chain 2 | Res2# | Res2 Name | Distance (Å) |
|---|---|---|---|---|---|---|---|---|
| 1 | 1-2 | B | 4 | GLY | Z | 8 | SER | 4.45 |
| 2 | 1-2 | B | 4 | GLY | Z | 9 | ASP | 4.04 |
| 3 | 1-2 | B | 29 | GLY | Z | 7 | THR | 3.89 |
| 4 | 1-2 | E | 12 | SER | P | 25 | TRP | 4.27 |
| 5 | 1-2 | E | 15 | GLU | P | 30 | ARG | 4.46 |
| 6 | 1-2 | G | 2 | ALA | H | 3 | GLU | 3.60 |
| 7 | 1-2 | G | 2 | ALA | a | 4 | GLY | 3.54 |
| 8 | 1-2 | G | 12 | SER | P | 8 | SER | 4.16 |
| 9 | 1-2 | L | 7 | THR | T | 29 | GLY | 4.40 |
| 10 | 1-2 | L | 15 | GLU | A | 4 | GLY | 3.86 |
| 11 | 1-2 | L | 17 | GLN | A | 2 | ALA | 3.99 |
| 12 | 1-2 | O | 2 | ALA | P | 4 | GLY | 4.34 |
| 13 | 1-2 | O | 4 | GLY | P | 4 | GLY | 4.41 |
| 14 | 1-2 | O | 10 | VAL | P | 1 | HIS | 3.86 |
| 15 | 1-2 | O | 27 | VAL | a | 10 | VAL | 4.12 |
| 16 | 1-2 | O | 28 | ARG | a | 7 | THR | 4.47 |
| 17 | 1-2 | b | 7 | THR | C | 31 | GLY | 4.22 |
| 18 | 1-2 | b | 8 | SER | C | 31 | GLY | 3.83 |

**(d)** 30-mer, 360 K, replicate 2

| Entry | Cluster Pair | Chain 1 | Res1# | Res1 Name | Chain 2 | Res2# | Res2 Name | Distance (Å) |
|---|---|---|---|---|---|---|---|---|
| 1 | 1-2 | C | 1 | HIS | I | 15 | GLU | 3.94 |
| 2 | 1-2 | C | 11 | SER | I | 10 | VAL | 4.34 |

| | | | | | | | | |
|---|---|---|---|---|---|---|---|---|
| 3 | 1-2 | C | 24 | ALA | a | 23 | ILE | 3.87 |
| 4 | 1-2 | J | 1 | HIS | I | 31 | GLY | 4.45 |
| 5 | 1-2 | J | 4 | GLY | I | 28 | ARG | 4.19 |
| 6 | 1-2 | J | 10 | VAL | a | 29 | GLY | 4.40 |
| 7 | 1-2 | Q | 7 | THR | W | 18 | ALA | 4.00 |
| 8 | 1-2 | Q | 20 | D6M | I | 6 | PHE | 3.51 |
| 9 | 1-2 | X | 24 | ALA | a | 14 | LEU | 3.93 |
| 10 | 1-2 | c | 1 | HIS | H | 11 | SER | 4.13 |
| 11 | 1-2 | c | 1 | HIS | H | 12 | SER | 3.84 |
| 12 | 1-2 | c | 2 | ALA | T | 18 | ALA | 3.95 |
| 13 | 1-2 | c | 3 | GLU | T | 30 | ARG | 3.53 |
| 14 | 1-2 | c | 5 | THR | H | 1 | HIS | 4.41 |
| 15 | 1-2 | c | 8 | SER | T | 15 | GLU | 3.64 |

**(e)** 45-mer, 300 K, replicate 1

| Entry | Cluster Pair | Chain 1 | Res1# | Res1 Name | Chain 2 | Res2# | Res2 Name | Distance (Å) |
|---|---|---|---|---|---|---|---|---|
| 1 | 1-2 | O | 29 | GLY | D | 15 | GLU | 3.85 |
| 2 | 1-2 | O | 29 | GLY | D | 16 | GLY | 3.82 |
| 3 | 1-2 | O | 31 | GLY | D | 17 | GLN | 4.30 |
| 4 | 1-2 | R | 14 | LEU | o | 10 | VAL | 4.37 |
| 5 | 1-2 | R | 16 | GLY | o | 9 | ASP | 3.99 |
| 6 | 1-2 | f | 20 | D6M | o | 10 | VAL | 4.40 |
| 7 | 1-2 | f | 27 | VAL | D | 2 | ALA | 4.25 |
| 8 | 1-2 | f | 28 | ARG | D | 2 | ALA | 4.10 |
| 9 | 1-2 | f | 31 | GLY | D | 2 | ALA | 4.22 |
| 10 | 1-2 | q | 1 | HIS | D | 18 | ALA | 4.10 |
| 11 | 1-2 | q | 9 | ASP | D | 18 | ALA | 4.25 |
| 12 | 2-3 | B | 3 | GLU | e | 4 | GLY | 4.27 |
| 13 | 2-3 | B | 9 | ASP | r | 7 | THR | 4.18 |
| 14 | 2-3 | B | 11 | SER | r | 7 | THR | 4.39 |

| | | | | | | | | |
|---|---|---|---|---|---|---|---|---|
| 15 | 2-3 | B | 11 | SER | r | 8 | SER | 4.02 |
| 16 | 2-3 | B | 12 | SER | r | 7 | THR | 4.48 |
| 17 | 2-3 | B | 19 | ALA | r | 23 | ILE | 4.27 |
| 18 | 2-3 | B | 25 | TRP | r | 24 | ALA | 3.84 |
| 19 | 2-3 | B | 31 | GLY | r | 16 | GLY | 3.86 |
| 20 | 2-3 | T | 24 | ALA | r | 1 | HIS | 3.49 |
| 21 | 2-3 | Z | 1 | HIS | e | 3 | GLU | 4.47 |
| 22 | 2-3 | Z | 3 | GLU | e | 2 | ALA | 3.71 |
| 23 | 2-3 | g | 1 | HIS | e | 9 | ASP | 4.38 |
| 24 | 2-3 | g | 18 | ALA | G | 19 | ALA | 4.44 |
| 25 | 2-3 | g | 25 | TRP | r | 27 | VAL | 4.38 |
| 26 | 2-3 | g | 26 | LEU | r | 28 | ARG | 4.11 |
| 27 | 2-3 | g | 27 | VAL | r | 29 | GLY | 4.43 |
| 28 | 3-4 | E | 5 | THR | c | 6 | PHE | 4.49 |
| 29 | 3-4 | E | 6 | PHE | U | 5 | THR | 4.32 |
| 30 | 3-4 | E | 7 | THR | U | 2 | ALA | 4.11 |
| 31 | 3-4 | E | 9 | ASP | U | 2 | ALA | 4.34 |
| 32 | 3-4 | E | 9 | ASP | U | 7 | THR | 3.93 |
| 33 | 3-4 | E | 15 | GLU | U | 1 | HIS | 3.83 |
| 34 | 3-4 | E | 23 | ILE | I | 31 | GLY | 4.45 |
| 35 | 3-4 | G | 5 | THR | U | 31 | GLY | 4.45 |
| 36 | 3-4 | G | 12 | SER | i | 2 | ALA | 3.88 |
| 37 | 3-4 | N | 1 | HIS | c | 11 | SER | 4.35 |
| 38 | 3-4 | N | 5 | THR | U | 4 | GLY | 3.69 |
| 39 | 3-4 | P | 10 | VAL | i | 18 | ALA | 4.12 |
| 40 | 3-4 | W | 1 | HIS | I | 18 | ALA | 3.93 |
| 41 | 3-4 | W | 2 | ALA | c | 5 | THR | 4.28 |
| 42 | 3-4 | W | 15 | GLU | I | 24 | ALA | 4.18 |
| 43 | 3-4 | W | 17 | GLN | c | 5 | THR | 4.44 |
| 44 | 3-4 | W | 18 | ALA | I | 25 | TRP | 4.48 |
| 45 | 3-4 | d | 17 | GLN | m | 19 | ALA | 4.01 |

| | | | | | | | | |
|---|---|---|---|---|---|---|---|---|
| 46 | 3-4 | d | 17 | GLN | m | 21 | GLU | 4.41 |
| 47 | 3-4 | d | 26 | LEU | I | 28 | ARG | 4.50 |
| 48 | 3-4 | d | 26 | LEU | I | 31 | GLY | 4.32 |
| 49 | 3-4 | d | 27 | VAL | I | 28 | ARG | 4.23 |
| 50 | 3-4 | d | 30 | ARG | I | 11 | SER | 4.43 |
| 51 | 3-4 | d | 31 | GLY | I | 14 | LEU | 4.48 |
| 52 | 3-4 | r | 14 | LEU | i | 18 | ALA | 4.19 |
| 53 | 3-4 | r | 18 | ALA | i | 8 | SER | 3.88 |
| 54 | 3-4 | r | 19 | ALA | i | 11 | SER | 4.41 |
| 55 | 3-4 | r | 19 | ALA | i | 12 | SER | 3.89 |

**(f)** 45-mer, 300 K, replicate 2

| Entry | Cluster Pair | Chain 1 | Res1# | Res1 Name | Chain 2 | Res2# | Res2 Name | Distance (Å) |
|---|---|---|---|---|---|---|---|---|
| 1 | 1-2 | F | 3 | GLU | q | 7 | THR | 4.37 |
| 2 | 1-2 | F | 9 | ASP | q | 4 | GLY | 4.45 |
| 3 | 1-2 | F | 10 | VAL | f | 8 | SER | 3.93 |
| 4 | 1-2 | F | 11 | SER | f | 9 | ASP | 4.15 |
| 5 | 1-2 | F | 11 | SER | q | 4 | GLY | 4.40 |
| 6 | 1-2 | F | 21 | GLU | q | 1 | HIS | 4.48 |
| 7 | 1-2 | I | 24 | ALA | M | 11 | SER | 3.78 |
| 8 | 1-2 | g | 1 | HIS | f | 4 | GLY | 4.15 |
| 9 | 1-2 | g | 10 | VAL | M | 20 | D6M | 4.18 |
| 10 | 1-2 | j | 2 | ALA | q | 11 | SER | 4.07 |
| 11 | 1-2 | j | 21 | GLU | M | 4 | GLY | 3.85 |
| 12 | 1-2 | j | 29 | GLY | q | 8 | SER | 4.07 |
| 13 | 2-3 | G | 3 | GLU | e | 8 | SER | 4.44 |
| 14 | 2-3 | G | 4 | GLY | e | 8 | SER | 3.37 |
| 15 | 2-3 | G | 4 | GLY | e | 9 | ASP | 4.11 |
| 16 | 2-3 | G | 24 | ALA | e | 1 | HIS | 4.07 |
| 17 | 2-3 | G | 31 | GLY | c | 19 | ALA | 4.25 |
| 18 | 2-3 | K | 4 | GLY | L | 4 | GLY | 4.49 |

| | | | | | | | | |
|---|---|---|---|---|---|---|---|---|
| 19 | 2-3 | K | 4 | GLY | L | 12 | SER | 3.90 |
| 20 | 2-3 | l | 3 | GLU | T | 1 | HIS | 4.04 |
| 21 | 2-3 | l | 4 | GLY | T | 2 | ALA | 4.38 |
| 22 | 2-4 | K | 22 | PHE | X | 10 | VAL | 3.98 |
| 23 | 2-4 | d | 2 | ALA | X | 2 | ALA | 4.27 |
| 24 | 2-4 | d | 5 | THR | X | 29 | GLY | 3.99 |
| 25 | 2-4 | d | 10 | VAL | X | 6 | PHE | 4.20 |
| 26 | 2-4 | l | 2 | ALA | X | 4 | GLY | 3.94 |
| 27 | 2-4 | l | 3 | GLU | X | 2 | ALA | 3.88 |
| 28 | 2-4 | l | 20 | D6M | X | 10 | VAL | 3.37 |
| 29 | 3-4 | c | 5 | THR | A | 30 | ARG | 3.93 |

**(g)** 45-mer, 360 K, replicate 1

| Entry | Cluster Pair | Chain 1 | Res1# | Res1 Name | Chain 2 | Res2# | Res2 Name | Distance (Å) |
|---|---|---|---|---|---|---|---|---|
| 1 | 1-2 | F | 31 | GLY | f | 4 | GLY | 4.36 |
| 2 | 1-2 | O | 6 | PHE | a | 31 | GLY | 4.41 |
| 3 | 1-2 | Z | 2 | ALA | P | 5 | THR | 4.12 |
| 4 | 1-2 | Z | 3 | GLU | P | 2 | ALA | 3.95 |
| 5 | 1-2 | Z | 12 | SER | V | 24 | ALA | 4.30 |
| 6 | 1-2 | Z | 13 | TYR | V | 23 | ILE | 4.49 |
| 7 | 1-2 | Z | 15 | GLU | V | 24 | ALA | 4.25 |
| 8 | 1-2 | Z | 19 | ALA | M | 8 | SER | 4.16 |
| 9 | 1-2 | e | 5 | THR | M | 19 | ALA | 4.45 |
| 10 | 1-2 | i | 17 | GLN | P | 2 | ALA | 4.40 |
| 11 | 1-3 | A | 1 | HIS | c | 15 | GLU | 4.42 |
| 12 | 1-3 | A | 1 | HIS | c | 18 | ALA | 3.64 |
| 13 | 1-3 | A | 17 | GLN | c | 17 | GLN | 4.10 |
| 14 | 1-3 | A | 17 | GLN | r | 4 | GLY | 4.26 |
| 15 | 1-3 | A | 18 | ALA | r | 7 | THR | 4.34 |
| 16 | 1-3 | A | 18 | ALA | r | 8 | SER | 3.61 |
| 17 | 1-3 | A | 19 | ALA | r | 4 | GLY | 4.39 |

| | | | | | | | | |
|---|---|---|---|---|---|---|---|---|
| 18 | 1-3 | A | 31 | GLY | c | 4 | GLY | 4.26 |
| 19 | 1-3 | B | 9 | ASP | c | 4 | GLY | 4.22 |
| 20 | 1-3 | B | 11 | SER | c | 5 | THR | 3.98 |
| 21 | 1-3 | B | 24 | ALA | c | 6 | PHE | 4.31 |
| 22 | 1-3 | B | 27 | VAL | L | 31 | GLY | 4.27 |
| 23 | 1-3 | B | 29 | GLY | L | 25 | TRP | 3.36 |
| 24 | 1-3 | O | 7 | THR | p | 5 | THR | 4.42 |
| 25 | 1-3 | O | 8 | SER | p | 9 | ASP | 4.43 |
| 26 | 1-3 | O | 11 | SER | H | 5 | THR | 4.27 |
| 27 | 1-3 | O | 22 | PHE | H | 4 | GLY | 4.46 |
| 28 | 1-3 | O | 22 | PHE | H | 24 | ALA | 3.90 |
| 29 | 1-3 | O | 29 | GLY | c | 9 | ASP | 3.84 |
| 30 | 1-3 | U | 4 | GLY | L | 3 | GLU | 3.56 |
| 31 | 1-3 | U | 4 | GLY | p | 19 | ALA | 4.36 |
| 32 | 1-3 | U | 5 | THR | p | 19 | ALA | 4.20 |
| 33 | 1-3 | U | 10 | VAL | r | 7 | THR | 4.06 |
| 34 | 1-3 | Z | 31 | GLY | N | 4 | GLY | 4.35 |
| 35 | 2-3 | M | 2 | ALA | m | 1 | HIS | 4.36 |
| 36 | 2-3 | M | 2 | ALA | m | 2 | ALA | 4.47 |
| 37 | 2-3 | M | 4 | GLY | m | 2 | ALA | 3.49 |
| 38 | 2-3 | M | 7 | THR | E | 15 | GLU | 3.98 |
| 39 | 2-3 | M | 7 | THR | E | 16 | GLY | 4.28 |
| 40 | 2-3 | V | 24 | ALA | K | 29 | GLY | 4.34 |
| 41 | 2-3 | a | 2 | ALA | K | 3 | GLU | 4.35 |
| 42 | 2-3 | a | 3 | GLU | m | 18 | ALA | 4.14 |
| 43 | 2-3 | a | 4 | GLY | K | 4 | GLY | 3.58 |
| 44 | 2-3 | a | 4 | GLY | m | 16 | GLY | 4.50 |
| 45 | 2-3 | a | 7 | THR | m | 25 | TRP | 4.11 |
| 46 | 2-3 | a | 10 | VAL | p | 29 | GLY | 3.71 |
| 47 | 2-3 | a | 18 | ALA | K | 7 | THR | 3.80 |
| 48 | 2-3 | a | 18 | ALA | K | 15 | GLU | 4.36 |

| | | | | | | | | |
|---|---|---|---|---|---|---|---|---|
| 49 | 2-3 | a | 19 | ALA | K | 15 | GLU | 3.81 |
| 50 | 2-3 | h | 17 | GLN | E | 30 | ARG | 4.10 |
| 51 | 2-3 | h | 22 | PHE | R | 24 | ALA | 3.75 |
| 52 | 2-3 | h | 24 | ALA | E | 19 | ALA | 4.00 |
| 53 | 2-3 | h | 24 | ALA | E | 21 | GLU | 3.98 |
| 54 | 2-3 | o | 3 | GLU | R | 2 | ALA | 3.57 |
| 55 | 2-3 | s | 8 | SER | K | 15 | GLU | 4.33 |

**(h)** 45-mer, 360 K, replicate 2

| Entry | Cluster Pair | Chain 1 | Res1# | Res1 Name | Chain 2 | Res2# | Res2 Name | Distance (Å) |
|---|---|---|---|---|---|---|---|---|
| 1 | 1-2 | F | 3 | GLU | e | 2 | ALA | 4.02 |
| 2 | 1-2 | F | 4 | GLY | e | 3 | GLU | 3.79 |
| 3 | 1-2 | P | 2 | ALA | N | 2 | ALA | 3.74 |
| 4 | 1-2 | P | 5 | THR | J | 4 | GLY | 4.05 |
| 5 | 1-2 | P | 5 | THR | J | 8 | SER | 3.68 |
| 6 | 1-2 | P | 7 | THR | J | 4 | GLY | 4.37 |
| 7 | 1-2 | P | 29 | GLY | J | 2 | ALA | 4.14 |
| 8 | 1-2 | P | 30 | ARG | p | 29 | GLY | 4.14 |
| 9 | 1-2 | V | 10 | VAL | e | 9 | ASP | 4.39 |
| 10 | 1-2 | h | 8 | SER | l | 29 | GLY | 4.31 |
| 11 | 1-2 | h | 9 | ASP | l | 4 | GLY | 4.14 |
| 12 | 1-2 | h | 17 | GLN | Y | 2 | ALA | 4.50 |
| 13 | 1-2 | h | 19 | ALA | e | 10 | VAL | 4.29 |
| 14 | 1-2 | h | 21 | GLU | e | 12 | SER | 4.18 |
| 15 | 1-2 | h | 25 | TRP | J | 6 | PHE | 4.41 |
| 16 | 1-2 | h | 29 | GLY | N | 9 | ASP | 3.95 |
| 17 | 1-2 | h | 31 | GLY | N | 1 | HIS | 3.89 |
| 18 | 1-2 | j | 1 | HIS | l | 7 | THR | 4.44 |
| 19 | 1-3 | C | 1 | HIS | i | 3 | GLU | 3.71 |
| 20 | 1-3 | C | 3 | GLU | A | 17 | GLN | 4.31 |

| | | | | | | | | |
|---|---|---|---|---|---|---|---|---|
| 21 | 1-3 | C | 5 | THR | A | 18 | ALA | 4.17 |
| 22 | 1-3 | C | 6 | PHE | A | 19 | ALA | 3.49 |
| 23 | 1-3 | M | 17 | GLN | H | 9 | ASP | 4.41 |
| 24 | 1-3 | Q | 1 | HIS | W | 12 | SER | 4.18 |
| 25 | 1-3 | S | 2 | ALA | a | 9 | ASP | 4.27 |
| 26 | 1-3 | S | 3 | GLU | H | 2 | ALA | 3.97 |
| 27 | 1-3 | S | 4 | GLY | H | 2 | ALA | 3.58 |
| 28 | 1-3 | S | 5 | THR | i | 19 | ALA | 4.18 |
| 29 | 1-3 | S | 21 | GLU | H | 1 | HIS | 4.44 |
| 30 | 1-3 | X | 4 | GLY | W | 24 | ALA | 4.49 |
| 31 | 1-3 | X | 4 | GLY | W | 28 | ARG | 4.24 |
| 32 | 1-3 | X | 6 | PHE | W | 20 | D6M | 4.01 |
| 33 | 1-3 | X | 11 | SER | i | 30 | ARG | 4.43 |
| 34 | 1-3 | X | 18 | ALA | i | 29 | GLY | 3.67 |
| 35 | 1-3 | X | 30 | ARG | i | 2 | ALA | 4.13 |
| 36 | 1-3 | c | 19 | ALA | H | 3 | GLU | 4.07 |
| 37 | 1-3 | c | 19 | ALA | H | 5 | THR | 3.84 |
| 38 | 1-3 | o | 3 | GLU | W | 8 | SER | 3.92 |
| 39 | 1-3 | o | 5 | THR | W | 6 | PHE | 4.08 |
| 40 | 1-3 | r | 5 | THR | i | 16 | GLY | 4.28 |
| 41 | 1-4 | F | 1 | HIS | G | 16 | GLY | 4.37 |
| 42 | 1-4 | F | 2 | ALA | G | 17 | GLN | 3.95 |
| 43 | 1-4 | I | 2 | ALA | B | 19 | ALA | 4.46 |
| 44 | 1-4 | I | 19 | ALA | O | 21 | GLU | 4.13 |
| 45 | 1-4 | L | 2 | ALA | B | 4 | GLY | 4.39 |
| 46 | 1-4 | L | 2 | ALA | G | 4 | GLY | 4.40 |
| 47 | 1-4 | L | 5 | THR | B | 2 | ALA | 4.41 |
| 48 | 1-4 | L | 5 | THR | B | 24 | ALA | 4.45 |
| 49 | 1-4 | L | 6 | PHE | B | 24 | ALA | 3.59 |
| 50 | 1-4 | L | 8 | SER | G | 7 | THR | 4.48 |
| 51 | 1-4 | L | 9 | ASP | G | 8 | SER | 3.93 |

| | | | | | | | | |
|---|---|---|---|---|---|---|---|---|
| 52 | 1-4 | S | 15 | GLU | B | 29 | GLY | 4.03 |
| 53 | 1-4 | S | 24 | ALA | B | 29 | GLY | 3.06 |
| 54 | 1-4 | S | 30 | ARG | B | 27 | VAL | 4.46 |
| 55 | 1-4 | S | 31 | GLY | B | 1 | HIS | 4.46 |
| 56 | 1-4 | j | 2 | ALA | U | 9 | ASP | 4.00 |
| 57 | 1-4 | j | 2 | ALA | U | 29 | GLY | 4.40 |
| 58 | 1-4 | j | 28 | ARG | O | 16 | GLY | 4.35 |
| 59 | 2-4 | K | 16 | GLY | q | 7 | THR | 3.76 |
| 60 | 2-4 | K | 21 | GLU | q | 4 | GLY | 4.29 |
| 61 | 2-4 | K | 25 | TRP | D | 11 | SER | 4.41 |
| 62 | 2-4 | K | 26 | LEU | D | 10 | VAL | 4.36 |
| 63 | 2-4 | Y | 8 | SER | G | 17 | GLN | 4.05 |
| 64 | 2-4 | n | 15 | GLU | U | 11 | SER | 4.46 |
